# Approaching the limit: Modeling the physiological bounds of playable space in football

**DOI:** 10.64898/2026.07.04.736520

**Authors:** Abdullah Zafar, Matthias Krüll, Samuel Guay, Louis De Beaumont

## Abstract

Pitch control models quantify spatial dominance in football by estimating which player can arrive first at each pitch location, but they treat all players as equivalently capable regardless of preceding effort. We introduce physiology-aware pitch control (Phys-PC), a model-agnostic time-to-arrive correction that imposes two physiological capacity channels calibrated from tracking data: a transient recoverable burden capturing incomplete recovery from recent high-intensity efforts, and a cumulative non-recoverable drain accumulating across match play. Both channels reduce a bounded access scale that modulates kinematic TTA before any downstream pitch-control computation. All parameters are anchored to exercise-physiology benchmarks; no laboratory measurements are assumed.

Applied to a 64-match international tournament, Phys-PC reveals structure that kinematic models cannot detect. In head-to-head races, the dominant burden channel shifts from transient to cumulative over the course of a match, with a transient resurgence in the final 15 minutes. These physiological asymmetries predict match outcomes: relative reserve advantage is associated with higher odds of winning ground challenges (OR = 1.20, *p* = 0.006; +4.2 pp), completing over-the-top passes past recovering defenders (OR = 1.45, *p* = 0.034; +8.3 pp), and progressing possession sequences into the final third (OR = 1.31, *p* = 0.013; +5.1 pp). At the player-profile level, an acute-cumulative decomposition of contested space access separates roles and individuals whose territorial reach is maintained through sustained positioning from those whose access is rebuilt through repeated high-intensity actions, providing a physiological lens on team tactical structure.

## 1 Introduction

Space is football’s most visible absence. It appears in the gaps between formations, representing both an opportunity to advance and a vulnerability to defend. However, these possibilities are only resolved through timely movement. Whether an attacker is running into the space or a defender is recovering to close it, outpacing the opponent is not merely a matter of kinematics; it is fundamentally limited by physical capacity, movement cost, and accumulated effort. Pitch control models quantify this spatial landscape by estimating which player can arrive first at each location on the pitch, transforming an abstract geometry into a continuous surface of territorial influence Spearman et al. [2017], Fernandez and Bornn [2018]. The foundational insight is that spatial dominance is determined not by where players stand but by how quickly they can move: time-to-arrive (TTA) under kinematic assumptions about acceleration and top speed drives the model, with each player implicitly claiming any region they can reach before opponents. Subsequent work has refined the underlying motion models Wu and Swartz [2025] and introduced player-specific top-speed variation da Silva et al. [2025]. Most notably, recent work has supplemented kinematic TTA with a scalar stamina factor that scales a player’s effective top speed, demonstrating the field’s movement toward capacity-aware control da Silva et al. [2025]. Such a scalar factor is useful for representing reduced movement capacity, but it cannot distinguish transient burden following high-intensity efforts from match-cumulative depletion that compounds across the full ninety minutes Bradley et al. [2010], Mohr et al. [2005].

The distinction matters because identical spatial configurations can impose different physical costs depending on the preceding passage of play. A fullback may occupy the same pitch location, but their capacity to close down a winger will differ if they arrive after a maximal recovery sprint rather than after a quiet minute. Exercise physiology provides a mechanistic basis for these effects: phosphocreatine (PCr) resynthesis after maximal sprinting occurs rapidly but remains incomplete over short inter-effort intervals, while high-intensity running output declines across match play in a manner consistent with cumulative fatigue Bogdanis et al. [1996], Bishop [2012], Bangsbo et al. [2006], Ørtenblad et al. [2013]. These mechanisms motivate the two capacity channels modeled here: transient recoverable burden and cumulative match-scale capacity drain. Existing pitch-control frameworks typically omit both mechanisms, treating reachability as a function of position, velocity, and fixed movement parameters. As a result, they may overestimate a player’s effective spatial reach when transient fatigue or cumulative load has altered the physical cost of arrival, particularly during late-match transitions, repeated pressing sequences, and defensive recoveries from back-to-back sprints.

We introduce physiology-aware pitch control (Phys-PC), a model-agnostic TTA correction that imposes tracking-derived physical-capacity constraints on any downstream pitch-control model. Existing kinematic pitch-control models estimate who can arrive first; physiology-aware pitch control asks whether that arrival remains physically affordable given the movement cost, transient fatigue, and cumulative load. The model encodes this through three channels computed from spatiotemporal tracking data: instantaneous relative metabolic demand, transient recoverable burden, and cumulative match-scale capacity drain. Together, these components modulate kinematic capacity before TTA is passed to the downstream pitch-control model, without altering the possession-weighting or control-surface logic. All parameters are calibrated from the match corpus, and no laboratory measurements are assumed.

The remainder of the paper is organized as follows. Section 2 describes the spatiotemporal tracking dataset and preprocessing pipeline. Section 3 presents the Phys-PC model and staged calibration pipeline. Section 4 reports component-level and combined-model validation, including face-validity probes for transient and cumulative capacity states across positional roles. Section 5 evaluates whether relative physiological reserve corresponds to observed match outcomes. Section 6 demonstrates three practical applications. Section 7 addresses model assumptions, limitations, and directions for extension.

## 2 Data

### 2.1 Spatiotemporal tracking and event data

The analysis uses proprietary tracking and event data from Pro Football Focus FC (PFF FC, Cincinnati, OH, USA), covering all 64 matches of the 2022 FIFA Men’s World Cup. Raw broadcast tracking is delivered by Sportlogiq (Montreal, QC, Canada); PFF applies a Kalman filter to all player location trajectories and smooths ball coordinates using event data, placing the ball at the carrier location and linearly interpolating between events. Both raw and Kalman-filtered coordinates are included in the data files; all physiological computations in this study use the smoothed player positions. Trajectories are recorded at 29.97 fps, providing continuous 2D pitch coordinates (*x, y*) in metres for all 22 outfield players and the ball per frame, synchronized with timestamped game and possession event annotations. Pitch dimensions are read from the per-match metadata (105 × 68 m across all 64 matches). The resulting data structure is equivalent to standard optical tracking datasets used in sports analytics Spearman et al. [2017]. Goalkeepers are excluded from all physiological calibration.

### 2.2 Data preprocessing and smoothing

No additional smoothing is applied beyond the provider-supplied Kalman filter, to avoid further signal attenuation. Instantaneous speed is estimated by lagged numerical differentiation of the smoothed positions over a 200 ms window, computed as the Euclidean displacement over that lag divided by the elapsed time. Forward acceleration is estimated from the lagged speed series using the same 200 ms window. Frame gaps exceeding 350 ms reset the motion state, preventing velocity contamination across stoppages or tracking interruptions. Because metabolic power (MP) is sensitive to high-frequency noise in optical tracking Savoia et al. [2020], the Kalman-filtered signal was validated to produce stable acceleration estimates prior to power computation.

Physiological calibration is restricted to full-match completers: outfield players present within five minutes of both kick-off and the final whistle, and meeting three tracking-reliability thresholds (stale-frame fraction ≤ 5%, clipped-acceleration fraction ≤ 5%, high-*q* fraction ≤ 5%). Players were grouped into five tactical roles: centre-backs (CB), fullbacks (FB), central midfielders (CM), wingers (W), and attacking midfielders/centre-forwards (AMCF). This criterion yields 1112 outfield player-match instances across all 64 matches (CB: 282, FB: 229, CM: 306, W: 136, AMCF: 159) and excludes players substituted for tactical or injury reasons whose load profiles would otherwise confound the maximum MP envelope and cumulative drain calibration.

## 3 Methods

### 3.1 Model overview

Physiological TTA (TTA_phys_) extends any kinematic TTA estimate in three steps: (1) estimate the player’s current physiological state from tracking history; (2) translate that state into a bounded effective access scale *ρ*; (3) compute TTA under kinematic caps scaled by *ρ*. The model has five equations and three latent states: normalized instantaneous demand *q*, transient recoverable burden *Z*, and cumulative non-recoverable drain *G* (Figure 1). All parameters are calibrated at the position-role level (CB, FB, CM, W, AMCF), reflecting meaningful between-role differences in sprint capacity, metabolic demand, and fatigue dynamics that remain interpretable above the smoothing noise of broadcast tracking.

**Figure 1.**
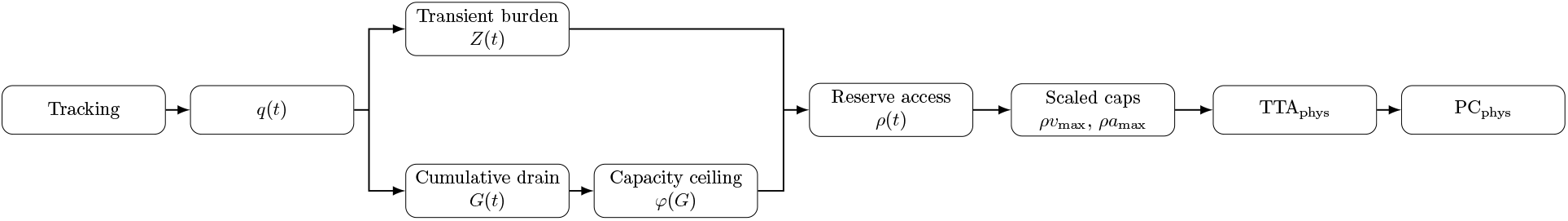
Model signal-flow diagram. Tracking inputs flow through normalized demand *q*(*t*) into two parallel latent-state paths: transient burden *Z*(*t*) and cumulative drain *G*(*t*). *G*(*t*) further modulates a capacity ceiling *φ*(*G*). Both paths feed the reserve access scale *ρ*(*t*), which scales kinematic caps (*ρv*_max_, *ρa*_max_) to produce TTA_phys_ and PC_phys_.

### 3.2 Metabolic demand and normalized intensity

Instantaneous metabolic power MP(*v, a*) is estimated using an equivalent-slope framework Savoia et al. [2020], di Prampero et al. [2005], Osgnach et al. [2010]. The equivalent-slope concept treats forward-accelerated running as biomechanically equivalent to uphill running at constant speed: the resultant of gravitational and forward acceleration defines an effective slope, which determines the energy cost per unit distance. A key property of this approach for football is that it captures the high energetic cost of acceleration at low and moderate speeds, where many decisive actions occur but conventional high-speed thresholds would assign near-zero load Osgnach et al. [2010].

Normalized demand *q* (Equation 1) expresses instantaneous intensity relative to the 95th-percentile MP expressed for that position role (MP95_ref_; calibrated in Section 3.9). It represents an intensity signal, not a fatigue state.

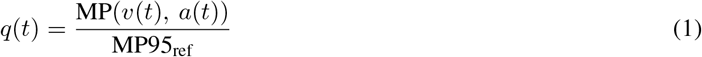

Because MP95_ref_ is role-specific, a CB and a winger performing the same sprint produce different *q* values, ensuring cross-role comparability. The equivalent-slope model is most reliable for straight-line accelerated running; its accuracy degrades for curvilinear paths, lateral shuffling, and change-of-direction actions Hader et al. [2016]. We therefore introduce an alignment penalty that reduces effective capacity for target-directed movements requiring substantial reorientation.

### 3.3 Kinematic movement profile and alignment penalty

For each target, the model evaluates candidate arrival times by constructing an accelerate-coast movement profile over the direct distance to that target. The profile is chosen to minimize metabolic cost while respecting player top speed and a global acceleration ceiling. Player-specific top speed *v*_max_ is estimated as the match-level p99.5 of smoothed speed, with per-role medians reported in Table 1. Acceleration is capped globally at *a*_max_ = 6.0m/s^2^, a literature-informed upper bound for elite soccer Yousefian et al. [2025]. We use a global acceleration cap rather than player-specific estimates because acceleration is less reliably recovered from broadcast tracking than top speed, and because a conservative ceiling prevents the profile optimizer from exploiting tracking noise.

**Table 1:** Per-role calibration results. *n*: outfield clean match completers. *τ* : LTI time constant derived by CV-3 burden-space minimisation. *Z*_crit_: transient burden threshold (p95, *q*-units). *v*_max_: median player top speed. *θ*_1_, *θ*_2_, *θ*_3_: CV-3 power-duration fit parameters.

| Role | $n$ | MP95<sub>ref</sub><br>(W/kg) | $\tau$<br>(s) | $Z_{\text{crit}}$<br>($q$-units) | $v_{\max}$<br>(m/s) | $\theta_1$<br>(W/kg) | $\theta_2$<br>(W s/kg) | $\theta_3$<br>(s) |
| --- | --- | --- | --- | --- | --- | --- | --- | --- |
| CB | 282 | 19.54 | 28.0 | 0.594 | 6.18 | 8.91 | 462.6 | 13.0 |
| FB | 229 | 21.37 | 28.0 | 0.613 | 6.61 | 9.40 | 473.2 | 12.2 |
| CM | 306 | 22.43 | 25.0 | 0.578 | 6.43 | 10.08 | 404.9 | 11.3 |
| W | 136 | 21.91 | 28.5 | 0.568 | 6.69 | 9.42 | 473.3 | 12.0 |
| AMCF | 159 | 21.88 | 26.5 | 0.579 | 6.42 | 9.45 | 425.9 | 11.4 |

The profile is computed along the direct ray to the target to isolate the physiological penalty from tactical path-planning choices. This amounts to a targeted counterfactual: given the same spatial trajectory, how do physiological constraints modulate arrival time? Rather than initializing from rest, the profile seeds the starting speed from the player’s current velocity projected onto the target direction, preventing artificial demand inflation for players already moving toward a location.

A player whose momentum is directed obliquely or away from the target cannot immediately transfer that speed; an alignment-penalized effective start speed is applied instead (Equation 2):

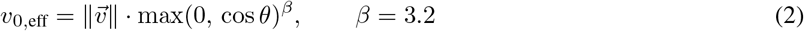

where *θ* is the angle between the player’s velocity vector and the target direction. The exponent *β* = 3.2 is calibrated to the empirical finding that a 45° change of direction redirects approximately 33% of forward momentum per execution step, penalising oblique approaches without modifying the standard metabolic equations Yamashita et al. [2025], Hader et al. [2016].

### 3.4 Power-duration capacity curve

Sustainable metabolic intensity is duration-dependent, as players can briefly exceed high relative power but cannot maintain it indefinitely. This power-duration (PD) relationship is well-established in exercise physiology via critical power theory Jones and Vanhatalo [2017], Poole et al. [2016] and has been demonstrated directly in soccer tracking data Mizelman et al. [2024]. Rather than importing laboratory-derived constants, the model employs an empirical per-role capacity curve *C*_PD_(*T*) fitted to rolling maximal metabolic power (MMP) envelopes derived from the tournament tracking data itself.

A three-parameter hyperbolic (CV-3) form is fitted per role (Equation 3):

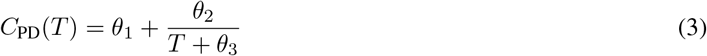

*θ*_1_ is the asymptotic sustainable capacity (the critical power analogue), *θ*_2_ scales the anaerobic reserve (the *W*′ analogue), and *θ*_3_ is a time-shift that stabilises the short-duration fit and prevents a singularity as *T* → 0 HUGH MORTON [1996]. Parameters are fitted per role by least-squares on the log-duration MMP envelope (Section 3.9). The capacity curve serves two distinct purposes in the model: it provides the MMP envelope against which the LTI time constant *τ* is calibrated (Section 3.9), and it defines the capacity ceiling that *φ*(*G*) progressively depresses across the match.

### 3.5 Transient burden state

A player’s physiological state seconds after a maximal sprint differs materially from that player at rest in the same location: phosphocreatine depletion, inorganic phosphate accumulation, and ionic disturbance all reduce contractile capacity over seconds-to-minutes timescales Bogdanis et al. [1996], Mendez-Villanueva et al. [2012]. Representing this transient incomplete-recovery dynamic requires an explicit memory state that tracks recent high-intensity load.

Transient burden *Z* is modelled as a first-order linear time-invariant (LTI) filter on normalised demand *q*, with a per-role decay constant *τ* (Equation 4):

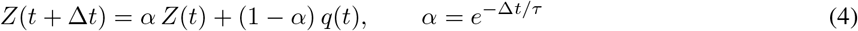

*Z* integrates recent high-*q* actions with exponential decay, with larger *τ* values indicating slower recovery and a longer burden memory. The per-role time constants (CB/FB: 28.0 s; CM: 25.0 s; W: 28.5 s; AMCF: 26.5 s) are calibrated from the empirical MMP envelope (Section 3.9) and are consistent with PCr resynthesis half-times and repeated-sprint recovery kinetics in the literature Bogdanis et al. [1996], McMahon and Jenkins [2002]. Thus, rather than representing a single metabolite, *Z* functions as a compressed proxy for rapidly reversible fatigue processes, including phosphocreatine depletion, metabolite accumulation, ionic disturbance, and glycolytic strain Girard et al. [2011]. Their aggregate recovery is approximated here by a single exponential decay.

*Z*/*Z*_crit_ is then defined as the per-role 95th percentile of *Z* across the full match corpus after calibrating *τ*. This provides a role-specific reference for high but routine transient burden, analogous to the MP95_ref_ normalization used for instantaneous demand. Transient utilisation is then expressed as *Z*/*Z*_crit_, so values near 1 indicate that a player is operating near the upper range of transient burden typically observed for that role.

### 3.6 Cumulative drain and capacity depression

Some components of match fatigue persist beyond short recovery intervals. Glycogen depletion, muscle damage, hyperthermia, and central fatigue can reduce late-match capacity despite brief low-intensity periods Mohr et al. [2005], Bangsbo et al. [2006], Ørtenblad et al. [2013]. We therefore distinguish the transient burden state *Z* from the cumulative capacity state *G*, modeled as monotone over the match.

Accumulated drain is translated into a capacity ceiling scalar *φ*(*G*) ∈ [*φ*_min_, 1] via Equation 5, which progressively depresses the power-duration curve over the match without altering instantaneous demand estimation, thereby preventing double-counting with the transient channel:

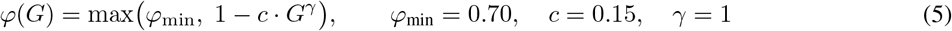

Cumulative drain *G* represents match-scale fatigue processes that do not substantially recover within a match. Unlike the transient burden state *Z*, which captures rapidly reversible limitations associated with PCr depletion and metabolite accumulation, *G* models the gradual loss of capacity arising from sustained energetic and mechanical stress. The monotone formulation is motivated by evidence that muscle glycogen is substantially depleted by half-time and is not meaningfully restored during the half-time interval Bangsbo et al. [2006], Krustrup et al. [2006], while repeated eccentric braking actions contribute structural muscle damage that accumulates across the full match Ørtenblad et al. [2013]. Accordingly, *G* is modeled as a cumulative state without within-match recovery.

The G channel operates on a separate normalisation from *Z*. Let

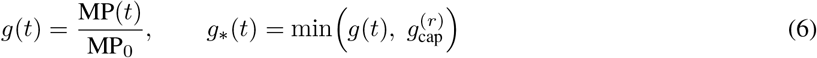

where MP_0_ = 14.96 W/kg is a fixed absolute metabolic-power threshold. Under this normalisation, *g* = 1 corresponds exactly to the onset of G-drain. Cumulative depletion then evolves according to

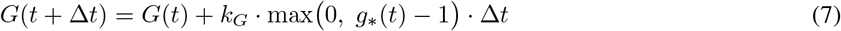

so only expenditure above the threshold contributes to long-term drain.

The capped input *g*_∗_(*t*) enforces the separation between the G and Z channels. The cap is defined as 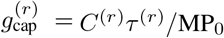, where *C*^(*r*)^*τ* ^(*r*)^ is the role-specific power-duration capacity evaluated at the transient LTI timescale. Expenditure beyond this limit is assumed to be predominantly PCr-driven and is therefore attributed to the transient burden state *Z* rather than to cumulative depletion. Across roles, 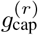 is approximately 1.40, corresponding to roughly 40% above the G-onset intensity.

### 3.7 Graded reserve access

Both *Z* and *G* reduce a bounded effective access scale *ρ* ∈ [*ρ*_min_, 1] that controls how much of a player’s remaining capacity the TTA extraction may draw on (Equation 8):

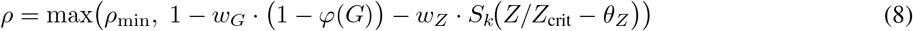

where 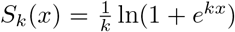 is the softplus function, smooth everywhere for finite *k* and converging to max(0, *x*) as *k* → ∞. Rather than vetoing arrival once a threshold is crossed, the bounded continuous scale reduces influence progressively with physiological burden. As a result, pitch-control surfaces remain continuous and all players continue to contribute at every physiological state.

The two penalty terms are structurally separated by design. The G-channel term *w*_*G*_ · (1 −*φ*(*G*)) reduces access in direct proportion to slow cumulative capacity loss, growing monotonically across the match. The transient channel term activates only once *Z* meaningfully exceeds the onset fraction *θ*_*Z*_ · *Z*_crit_ via the softplus gate, keeping the two mechanisms distinct. The floor is set to *ρ*_min_ = *φ*_min_ = 0.70, matching the capacity floor of the G-drain ceiling function so that the access scale cannot undercut the capacity level already established by *φ*(*G*).

The four penalty parameters are anchored to physiological benchmarks rather than optimised freely. At full G-drain (*φ* = *φ*_min_ = 0.70, so 1 − *φ* = 0.30), *w*_*G*_ = 0.50 yields a maximum kinematic reduction of 0.50 × 0.30 = 0.15, matching the 10-15% late-match high-intensity running decline documented in elite football Bradley et al. [2010]. At repeated-sprint ability (RSA) load (3 × 30 m sprints, 15 s passive rest), *w*_*Z*_ = 0.35 produces an ≈8% reduction in access, consistent with the 5-8% sprint-time decrements observed in repeated-sprint protocols Bishop [2012], Girard et al. [2011]. The onset *θ*_*Z*_ = 0.70 is set so that the same RSA protocol places all roles above the softplus midpoint 15 s after the final sprint, while an isolated sprint followed by ≥45 s recovery falls well below it. The sharpness parameter *k* = 10 ensures this penalty is applied smoothly rather than as a hard threshold. It distributes the 10-90% activation range of 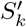 across a window of ≈0.44 normalised units (*Z*/*Z*_crit_ ∈ [0.50, 0.94]). Physiologically, this maps directly to the 15-30 s inter-sprint rest interval where a player’s capacity transitions rapidly from heavily impaired to substantially recovered Bishop [2012], Girard et al. [2011].

### 3.8 Physiology-adjusted TTA

The access scale *ρ* enters the TTA computation by scaling both kinematic caps simultaneously: *v*_eff_ = *v*_max_ · *ρ* and *a*_eff_ = *a*_max_ · *ρ*. TTA_phys_ is then the first candidate time *T* at which the accel-coast movement profile under (*v*_eff_, *a*_eff_) satisfies kinematic feasibility, found by a discrete scan over candidate times at 0.05 s resolution. The scan ceiling of 24 s is set conservatively: even at the minimum access floor (*ρ* = 0.70) applied to a typical outfield top speed, the resulting effective speed is sufficient to reach 40 m well within the scan window, so no physiologically relevant arrival is censored. The resulting TTA_phys_ replaces the kinematic TTA passed to any downstream pitch-control integration; no modification to the possession-weighting or control-surface logic is required.

### 3.9 Calibration pipeline

All physiological parameters are calibrated before any downstream analysis. The pipeline is strictly staged: each stage consumes only outputs of prior stages with no circular dependencies, and all parameters are frozen after calibration and held fixed for the validation and application analyses (Figure 2).

**Figure 2.**
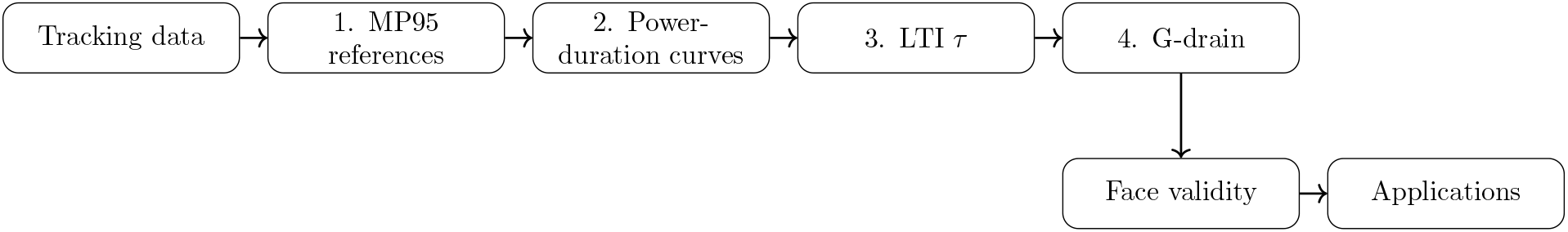
Calibration dependency graph. The pipeline is strictly staged with tracking data feeding four sequential calibration stages (MP95 references → power-duration curves → LTI *τ* → G-drain), after which all parameters are frozen. Final outputs feed a face-validity check before downstream applications.

#### 3.9.1 Stage 1: MP95 profiles

The reference intensity ceiling MP95_ref_ anchors the normalized demand signal *q*. It is estimated as the tournament-level median of each player’s match-level 95th-percentile metabolic power, then pooled within role. Extracted role-level values are reported in Table 1.

#### 3.9.2 Stage 2: Power-duration curves

The role-level maximal metabolic power envelope captures what a player in each role can sustain for any given duration. At each of 10 durations spanning 1 to 300 s, we take the p99.5 of the per-player rolling MMP distribution, pooled within role. The p99.5 gives a robust population ceiling that reflects genuine physiological capacity without being distorted by single-match outliers or tactical outliers. A three-parameter hyperbolic curve is then fit per role by least-squares on the log-duration envelope. The goodness-of-fit and literature anchor comparisons are reported in Section 4. Per-role parameters (*θ*_1_, *θ*_2_, *θ*_3_) are listed in Table 1.

#### 3.9.3 Stage 3: LTI time-constant derivation

The recovery time-constant *τ* controls how quickly the transient burden *Z* decays between efforts. Rather than assuming a value from the literature, we derive *τ* directly from the shape of the fitted MMP curve. The core idea is that if *τ* correctly captures the underlying recovery timescale, maximal efforts of different durations should accumulate similar levels of transient burden. Accordingly, a 5-second sprint and a 60-second hard run would impose comparable burdens on a properly calibrated filter. Formally, we define the burden implied by the PD curve as:

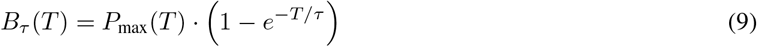

and search for the *τ* that minimises the coefficient of variation of *B*_*τ*_ (*T*) over *T* ∈ [10, 180] s. The derived values (CB/FB: 28.0 s; CM: 25.0 s; W: 28.5 s; AMCF: 26.5 s) fall within the PCr resynthesis literature band of 27-32 s for trained athletes Bogdanis et al. [1996], McMahon and Jenkins [2002]. The CV of *B*_*τ*_ (*T*) drops from 0.26-0.29 (raw MMP-duration) to 0.05-0.09 after derivation, confirming a well-defined minimum and validating the filter structure. Per-role *τ* and *Z*_crit_ are reported in Table 1.

#### 3.9.4 Stage 4: G-drain calibration

G-drain calibration sets the rate at which cumulative high-intensity work depletes match-scale capacity. The shape parameters are fixed by design: *c* = 0.15 and *γ* = 1, so that *G* = 1 represents full match-scale depletion with a 15% capacity loss (*φ*(1) = 0.85). The remaining free parameter, the drain rate *k*_*G*_, is anchored to a single empirical quantity: the player at the 75th percentile of tournament cumulative high-intensity load should reach *G* = 1 by match end. Concretely, *k*_*G*_ = 1/*L*_*ref*_, where *L*_*ref*_ is the p75 of 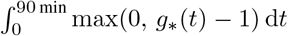 computed from tracking kinematics across match-completing outfield players. The derived value is *k*_*G*_ = 0.00469 (*L*_ref_ = 213.3 s). The resulting match-progression decline in high-MP output is shown in Figure 3 as a face-validity diagnostic rather than a calibration target.

**Figure 3.**
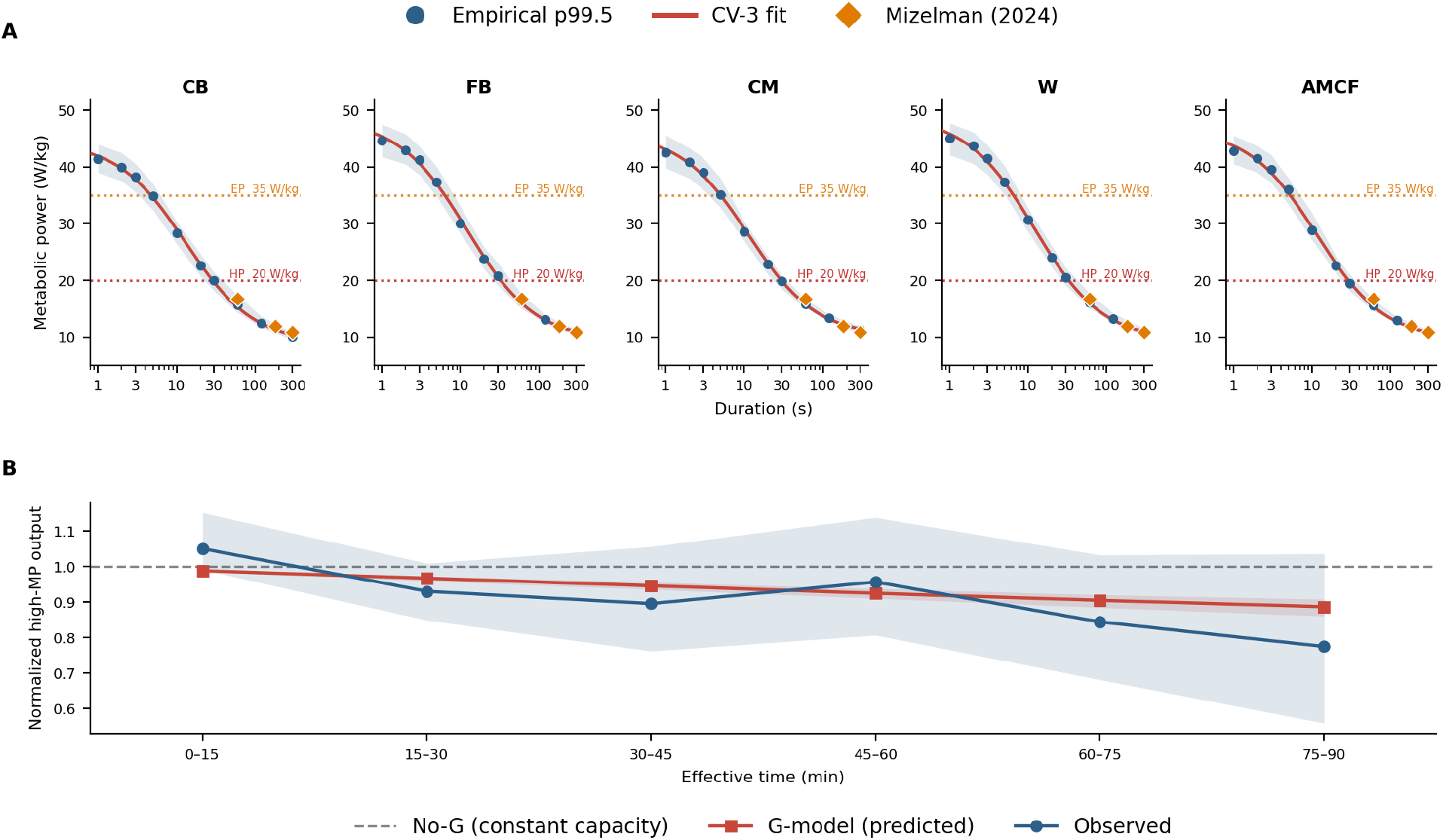
PD-curve and cumulative burden parameter validation. **A** Per-role CV-3 power-duration fits (red line) overlaid on the empirical tournament MMP envelope (blue points; shading = match-level IQR). Orange diamonds are independent literature anchors converted to metabolic power at 60, 180, and 300 s Mizelman et al. [2024]. Dotted horizontal lines mark the high-power (HP, 20 W/kg) and elevated-power (EP, 35 W/kg) zone boundaries Akyildiz et al. [2022]. All five roles share the same *y*-axis; the CB ceiling is visibly lower than the CM and W ceilings. **B** Block-level normalized high-MP output across six 15-minute effective-time blocks: observed (blue; shading = IQR), G-model predicted capacity scale *φ*(*G*) (red), and no-G constant-capacity baseline (dashed grey). The G-model drops only to *φ* 0.89 because the anchor is set at the p75 load player; the median player accumulates *G* ≈ 0.76 by match end, not full depletion. The G-model RMSE (0.064) is less than half the no-G RMSE (0.126), confirming the directional late-match decline is captured without early-match deflection.

## 4 Results

### 4.1 Power-duration fit

The tournament-derived MMP envelope is compared against published speed-duration and metabolic-power anchors from elite soccer to confirm that the model’s structural upper bounds are physiologically plausible. Key anchor points are: outfield p99.5 at 1 s (≈43.2 W/kg), 10 s (≈29.3 W/kg), and 60 s (≈15.8 W/kg); these fall within ranges reported for elite outfield players in Mizelman et al. [2024] and are consistent with critical-power analogues from running physiology Jones et al. [2010], Poole et al. [2016]. Per-role CV-3 fits overlaid on the empirical MMP envelope are shown in Figure 3.

### 4.2 LTI time-constant derivation

The derived *τ* values are compared against the literature RSA/VO_2_ kinetics band of 27-32 s and the match-level IQR of 24.5-28.0 s Bogdanis et al. [1996]. All five roles fall within or immediately adjacent to the literature band, confirming that the population MMP envelopes encode a recovery timescale consistent with PCr resynthesis kinetics in trained athletes McMahon and Jenkins [2002].

**Table 2:**
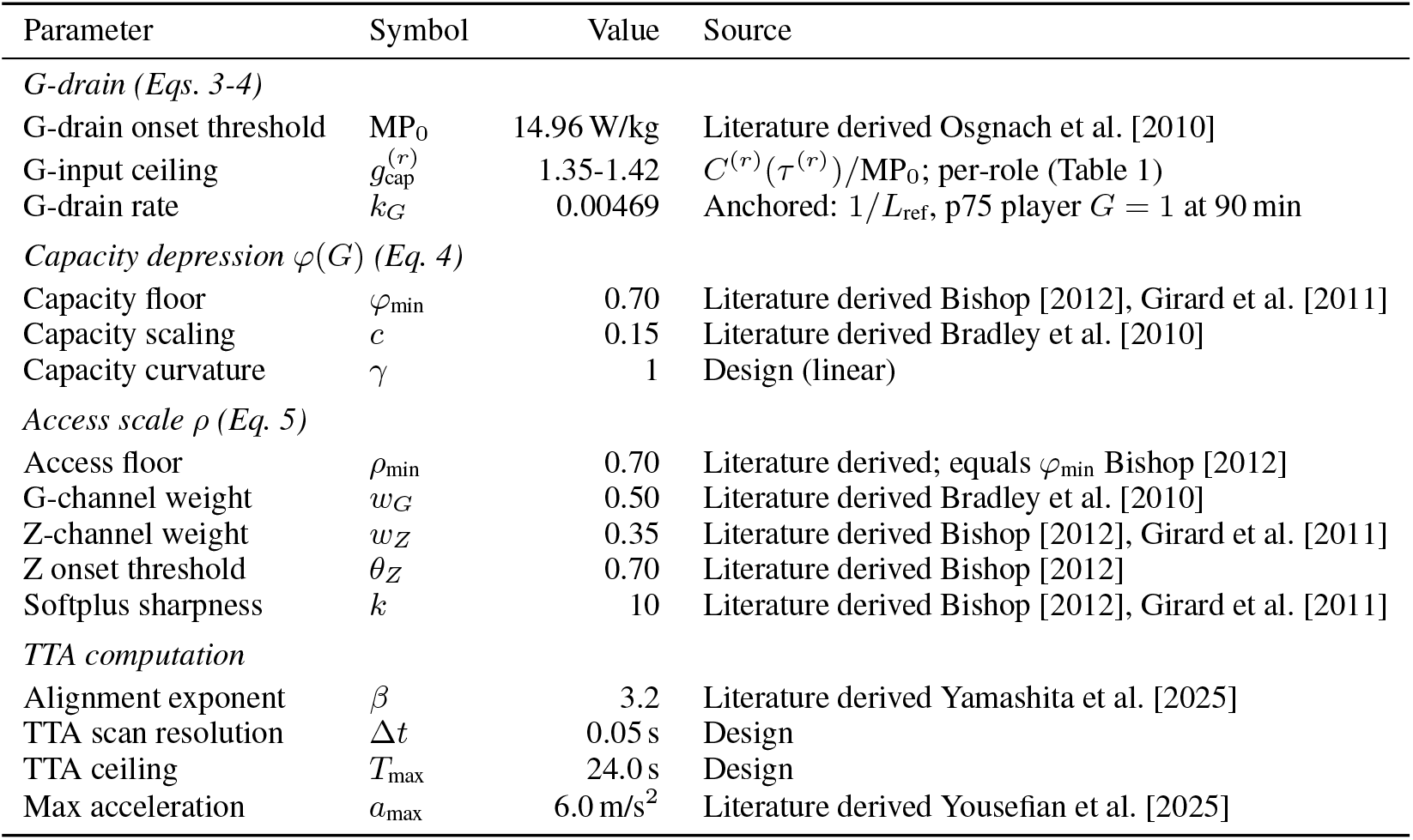
Full model parameter summary. Per-role parameters (MP95_ref_, *τ*, *Z*_crit_, *v*_max_, PD *θ*_1_-*θ*_3_) are reported in Table 1.

### 4.3 G-drain calibration fit

Observed versus predicted 6-block high-MP decline is shown as a face-validity diagnostic for the anchored *k*_*G*_. The no-G RMSE is 0.126 (constant capacity); the G model RMSE is 0.064. The primary check is qualitative: the model should reproduce the directional late-match decline without early-match deflection. Observed high-MP output versus predicted across the six effective-time 15-minute blocks is shown in Figure 3.

### 4.4 Combined model face validity

With all parameters frozen, the fully integrated model is probed for structural coherence across three complementary diagnostics (Figure 4).

**Figure 4.**
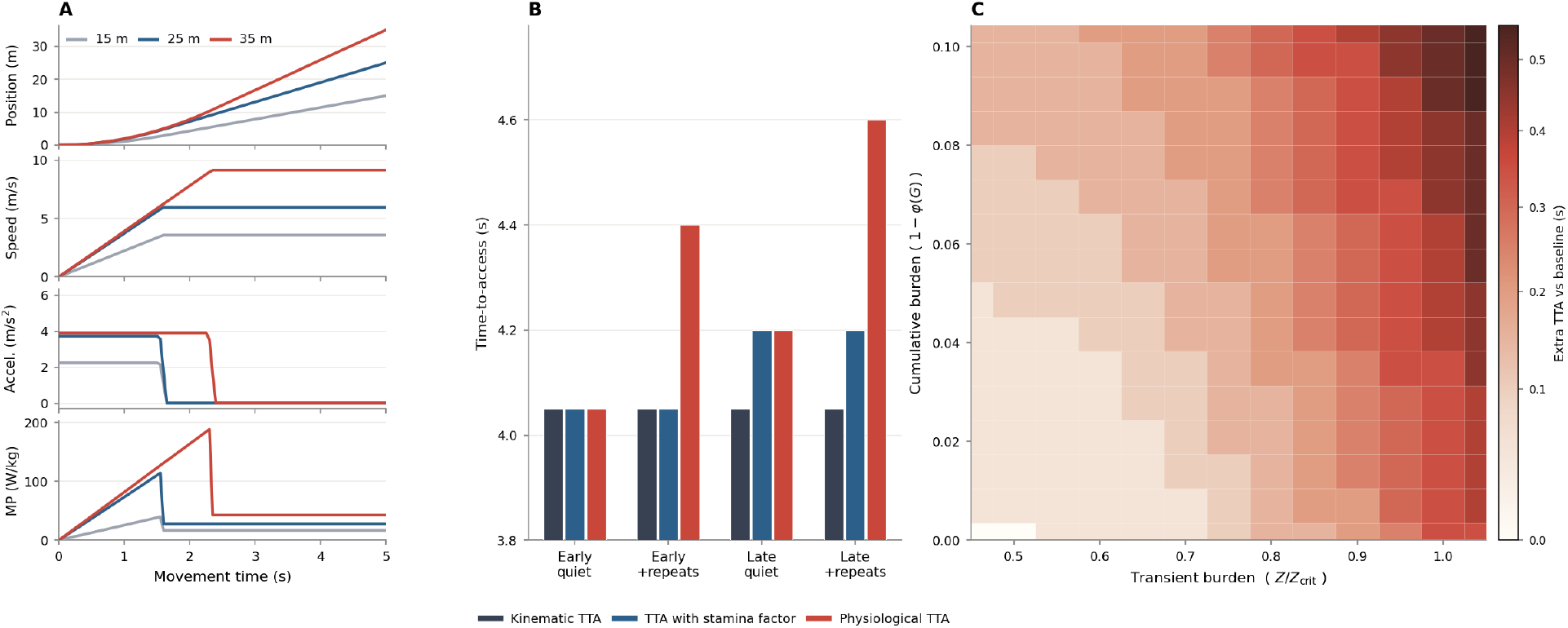
Face validity. **A:** Minimum-cost speed profiles for three target distances (15, 25, 35 m) at a fixed 5-second time budget from rest. Peak speed and initial acceleration increase with distance. **B:** Time-to-access (30 m probe) for three model variants across four canonical physiological states. Kinematic TTA (dark grey) is invariant to physiological state. TTA with a stamina factor (blue) captures the late-match cumulative G-drain effect. Physiological TTA (red) responds to both cumulative (G) and transient (Z) channels. **C:** Physiological extra TTA (s) response surface as a function of transient load and cumulative capacity loss. Colour scale from white (no extra TTA) to dark red (maximum penalty).

**Panel A** verifies the path optimiser. At a fixed 5-second time budget, minimum-cost-temporal profiles are computed for three target distances (15, 25, and 35 m) starting from rest. Longer target distances demand higher peak speeds and steeper initial acceleration, confirming that the optimiser trades off metabolic cost against deadline in a physically sensible manner.

**Panel B** isolates the physiological state-sensitivity of each model variant across four canonical match states: early-match quiet (low *Z, G* ≈ 0), early-match after repeated sprint efforts (high *Z, G* ≈ 0), late-match quiet (low *Z*, high *G*), and late-match after repeated efforts (high *Z*, high *G*). The probe uses five position roles as the source of variation; each role uses its own calibrated *τ*, *Z*_crit_, PD curve, and MP95_ref_, while kinematics are standardised (*v*_0_ = 3.0 m/s, top speed = 9.5 m/s) so that role differences reflect physiology only. Three model variants are compared: player-v_max_ TTA (state-invariant kinematic baseline), scalar-G TTA (cumulative drain only, no role spread), and full-reserve TTA (both channels with role-specific differentiation). As expected, player-v_max_ TTA is flat across all states and roles; scalar-G TTA steps up for late-match states but produces no role spread; full-reserve TTA differentiates all four states with role-specific transient burden trajectories, recovering toward the G-only value in the quiet late state as *Z* dissipates.

**Panel C** shows the joint response surface of full-reserve TTA as a function of transient load utilisation (*Z/Z*_crit_) and cumulative capacity loss (1 − *φ*(*G*)), evaluated at a standardised 30-m sprint. Extra TTA relative to a fresh baseline increases monotonically with both dimensions. The surface is smooth and the transition at the softplus threshold (*θ*_*Z*_ = 0.70) is continuous, confirming that no kinematic discontinuities are introduced.

## 5 Outcome validation across tactical scales

The preceding analyses show that Phys-PC produces physiologically plausible state dynamics. The next step is to determine whether these pre-action states carry information that is relevant to match outcomes. We therefore evaluated three event families ordered from micro to macro scale: non-aerial ground challenges, over-the-top passes, and territorial progression by possession sequences. These were chosen because each has a clear outcome, a plausible pre-event physiological mechanism, and an identifiable opponent or defensive unit contesting the action.

The key predictor in each analysis was a *relative* reserve advantage, not an actor’s absolute *ρ* alone. This is the main operational interpretation of Phys-PC: it does not claim that a fresh player will always succeed, but identifies when an opponent-relative physiological gap is opening or closing. That gap is not directly represented in conventional event-based or workload-based analyses because it depends on the interaction between recent movement history, calibrated reserve capacity, and the specific opponent contesting the action. All Phys-PC parameters were calibrated from tracking-derived power-duration curves and fixed before outcome analysis. We then fit binomial generalized linear mixed models with match-level random intercepts and event-family-specific controls Bates et al. [2015]. Because the models were fit using logistic regression, effect sizes are presented as odds ratios in Figure 5. For interpretability, Table 3 also reports the corresponding adjusted differences on the probability scale as percentage-point changes in success rate.

**Table 3:** Outcome-validation model summaries. Odds ratios compare Q4 versus Q1 relative-reserve advantage. Adjusted percentage-point differences are marginal fitted-rate translations from the same GLMMs.

| Outcome | Relative reserve contrast | Model $n$ | OR (95% CI) | $p$ | Adj. $\Delta$ |
| --- | --- | --- | --- | --- | --- |
| Ground-challenge win | Actor-opponent, 2 s pre-event | 8,109 | 1.20 (1.05-1.37) | 0.006 | +4.16 pp |
| Over-the-top completion | Target-nearest defender, 5 s pre-release | 1,275 | 1.45 (1.03-2.04) | 0.034 | +8.33 pp |
| Final-third entry | Middle-third unit gap at sequence start | 3,755 | 1.31 (1.06-1.61) | 0.013 | +5.06 pp |

**Figure 5.**
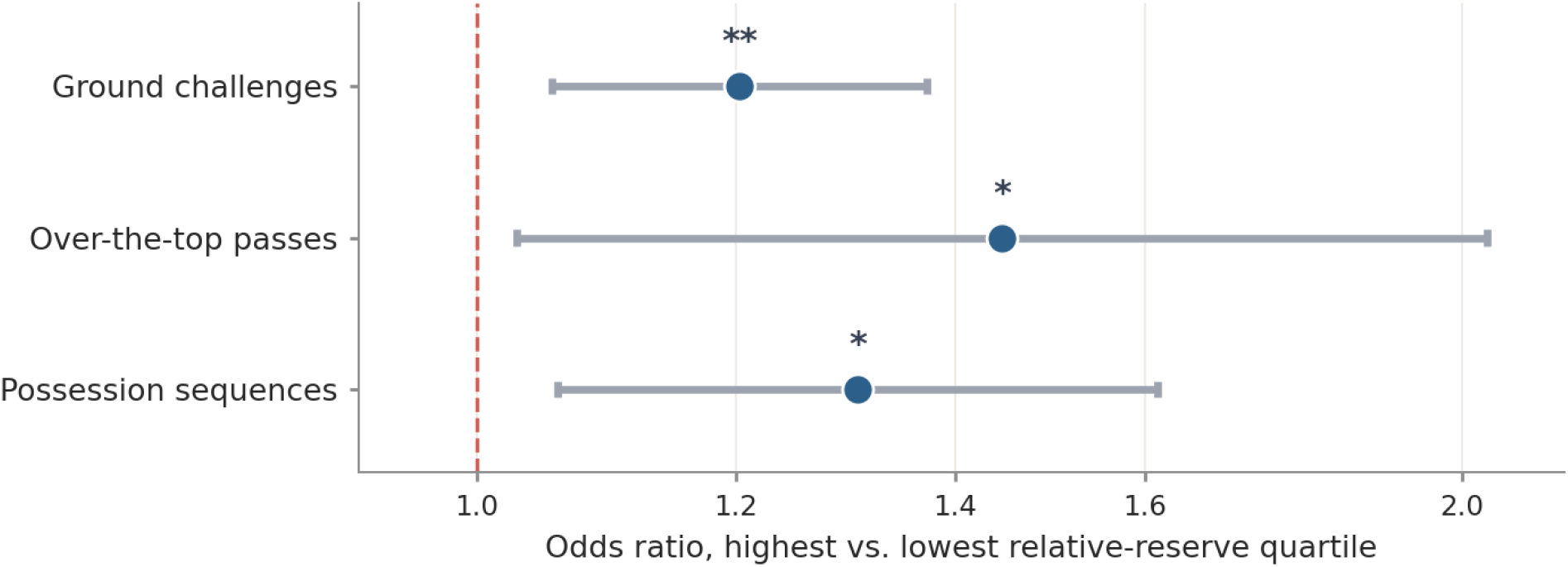
Validation against match outcomes. Odds ratios compare the highest versus lowest relative-reserve quartile from event-family-specific GLMMs with contextual controls and match-level random intercepts. Horizontal intervals show approximate 95% confidence intervals; the dashed line marks no association (OR=1). Significance: * *p* < 0.05, ** *p* < 0.01.

To benchmark the Phys-PC outcome models against simpler alternatives, we also fit matched tactical-context and scalar-stamina specifications on the same outcome samples. The tactical-context model retained only the event-family-specific controls and match random intercept. The scalar-stamina model added a quartile of the cumulative capacity-scale gap, computed from *φ*(*G*) for the same actor-opponent or team-unit contrast used in the Phys-PC analysis. These baselines were then compared with the Phys-PC specification described above, which used the corresponding relative-reserve quartile based on *ρ*. Thus, the specifications differed only in whether the tactical-context model was left unaugmented, augmented with a scalar cumulative-stamina contrast, or augmented with the full Phys-PC relative-reserve contrast. Likelihood-ratio tests compare each augmented model against the tactical-context specification.

### 5.1 Ground-challenge success

Ground challenges provide the most local test of the relative-advantage mechanism. Duels are among the few match actions where two players physically contest the same location at the same moment, making them a direct proxy for the capacity to arrive first, apply force, and sustain body position under contact. Because the outcome depends on acceleration, balance, and reactive agility over a timescale of 1-2 seconds, a player whose transient reserve is depleted from a preceding sprint or pressing sequence is plausibly less able to win the contest, even if their nominal kinematic profile suggests otherwise.

For non-aerial PFF challenge events (*n* = 8,109), we sampled player states 2 s before the action and compared the actor’s *ρ* with the opponent’s *ρ*. The highest reserve-advantage quartile won at 20% higher odds than the lowest (OR = 1.20, 95% CI 1.05-1.37, *p* = 0.006; adjusted Δ = +4.16 pp). The quartile pattern was positive but less monotonic than in the other analyses, with the largest fitted win rate in Q3. We therefore interpret this result as evidence that reserve asymmetry contributes to ground-contest outcomes, while acknowledging that challenge success is also strongly shaped by event geometry and contest type.

### 5.2 Over-the-top pass completion

Over-the-top passes extend the test to a race-to-space action where the physical contest between runner and recovering defender is the primary determinant of success. Unlike short passes, which are resolved largely by technique and positioning, an over-the-top ball bypasses midfield pressure entirely and converts the action into a footrace behind the defensive line. This makes over-the-top passes a high-leverage tactical tool: they force the opposition to choose between maintaining a high defensive line and risking exposure to depth, or dropping deep and conceding midfield territory. As a result, over-the-top passes are among the pass types most directly sensitive to runner-defender physiological asymmetry, because the outcome depends on who can close the distance to the landing zone first.

Here the relevant contest is not the passer in isolation, but the intended runner against the defender most able to recover the target space. We therefore sampled player states 5 s before release and used the target runner’s *ρ* minus the nearest defender’s *ρ* as the Phys-PC predictor (*n* = 1,275). The highest runner-advantage quartile completed at 45% higher odds than the lowest (OR = 1.45, 95% CI 1.03-2.04, *p* = 0.034; adjusted Δ = +8.33 pp), the largest effect among the three outcome families. This result illustrates the practical value of the relative metric: it identifies when a runner-defender physiological gap exists before the pass is released, rather than only describing the pass after completion or failure.

### 5.3 Possession-sequence progression

Possession sequences extend the mechanism to the unit level. Territorial progression through the middle third tests whether aggregate physiological balance, rather than a single player’s state, shapes collective spatial outcomes. The middle third is the contested zone where possession sequences are won or lost: the team that can press, support, and transition more effectively in this region controls whether the ball advances or recycles. When one side holds a physiological reserve advantage across multiple players in this zone, they can sustain pressing intensity, close passing lanes, and support forward runs at a higher collective tempo.

Because these sequences originate in the middle third, we quantified midfield reserve balance at sequence onset by comparing the mean *ρ* of possessing-team outfield players located in the middle third with the mean *ρ* of defending-team outfield players in the same zone (*n* = 3,755). This measure is not intended to describe the full possession trajectory; rather, it asks whether the physiological balance in the midfield contest at possession onset corresponds to subsequent territorial progression. Higher possessing-team reserve advantage was associated with 31% higher odds of entering the final third (OR = 1.31, 95% CI 1.06-1.61, *p* = 0.013; adjusted Δ = +5.06 pp), with a positive ordered-quartile trend (Figure 5; Table 3).

The comparison indicates that the Phys-PC relative-reserve quartile contributed information beyond tactical context for the two most direct contest outcomes: ground challenges and over-the-top passes (Table 4). The scalar cumulative-stamina baseline did not improve fit for either outcome. For possession-sequence progression, both scalar stamina and Phys-PC modestly improved fit relative to tactical context, though Phys-PC produced the stronger Q4-versus-Q1 contrast.

**Table 4:**
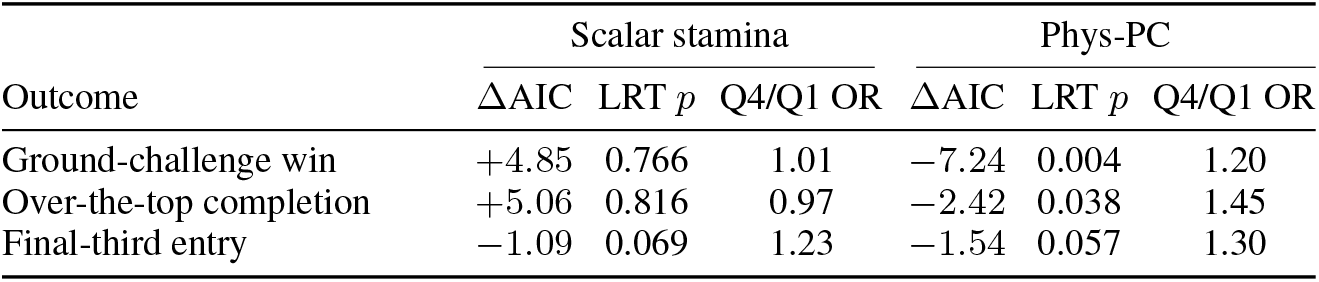
Incremental comparison against tactical-context and scalar-stamina baselines. Both augmented models use the same outcome-specific controls and match random intercepts as the tactical-context model. Scalar stamina is based on the cumulative capacity-scale gap *φ*(*G*); Phys-PC is based on the relative reserve gap *ρ*. ΔAIC is relative to the tactical-context model, and likelihood-ratio tests compare each augmented model with that tactical baseline.

| Outcome | Scalar stamina |  |  | Phys-PC |  |  |
| --- | --- | --- | --- | --- | --- | --- |
| | $\Delta\text{AIC}$ | LRT $p$ | Q4/Q1 OR | $\Delta\text{AIC}$ | LRT $p$ | Q4/Q1 OR |
| Ground-challenge win | +4.85 | 0.766 | 1.01 | −7.24 | 0.004 | 1.20 |
| Over-the-top completion | +5.06 | 0.816 | 0.97 | −2.42 | 0.038 | 1.45 |
| Final-third entry | −1.09 | 0.069 | 1.23 | −1.54 | 0.057 | 1.30 |

## 6 Applications

All parameters are frozen from calibration; no further tuning is applied. The following three applications demonstrate the practical utility of Phys-PC on events extracted from the full 64-match tournament dataset.

### 6.1 Head-to-head race

#### 6.1.1 Motivation

A head-to-head race where two players converge on the same open location is the canonical reachability scenario for pitch-control models. Kinematic pitch control is well-specified for this case when players are physiologically fresh. The transient *Z* channel captures within-possession burden from sprint sequences and repeated defensive efforts, and races anchored immediately after those efforts are where kinematic and physiological TTA are most likely to diverge in a structured way.

#### 6.1.2 Event extraction

Race candidates are anchored to four event types (pass sequence completions, clearances, contested backward passes, and receptions) that share a common structure: a target location in open space with at least one player from each team converging toward it. Candidates are filtered to retain spatially open 1-v-1 contests in which both players actively close toward the target, with a final proximity filter retaining only close contests where the kinematic TTA gap is small enough that physiological state could plausibly shift the model’s assessment.

#### 6.1.3 Findings

At each race anchor frame, both players’ full physiological state (*Z, G, φ*(*G*), *ρ*) is read from the match cache and TTA_phys_ is computed for each contestant. Events where the kinematic and physiological TTA rankings disagree are cases where accounting for physiology changes which player the model assigns the access advantage to.

Across the 64-match tournament the dominant channel shifts non-monotonically with match time (Figure 6A). Transient-driven shifts in PC predominate through the first half and into the early second half. From approximately the 60th minute onward, cumulative-driven shifts become the majority: this period coincides with the typical timing of substitutions, which introduce players with near-zero *G* accumulation against starters carrying substantial load, creating discrete G gaps that the kinematic model cannot see. Notably, this shift is not sustained: in the final 15 minutes, as teams push higher and the game opens up, the frequency of transient-dominant shifts rises again, reflecting sprint-intensive pressing sequences that elevate *Z* burden.

**Figure 6.**
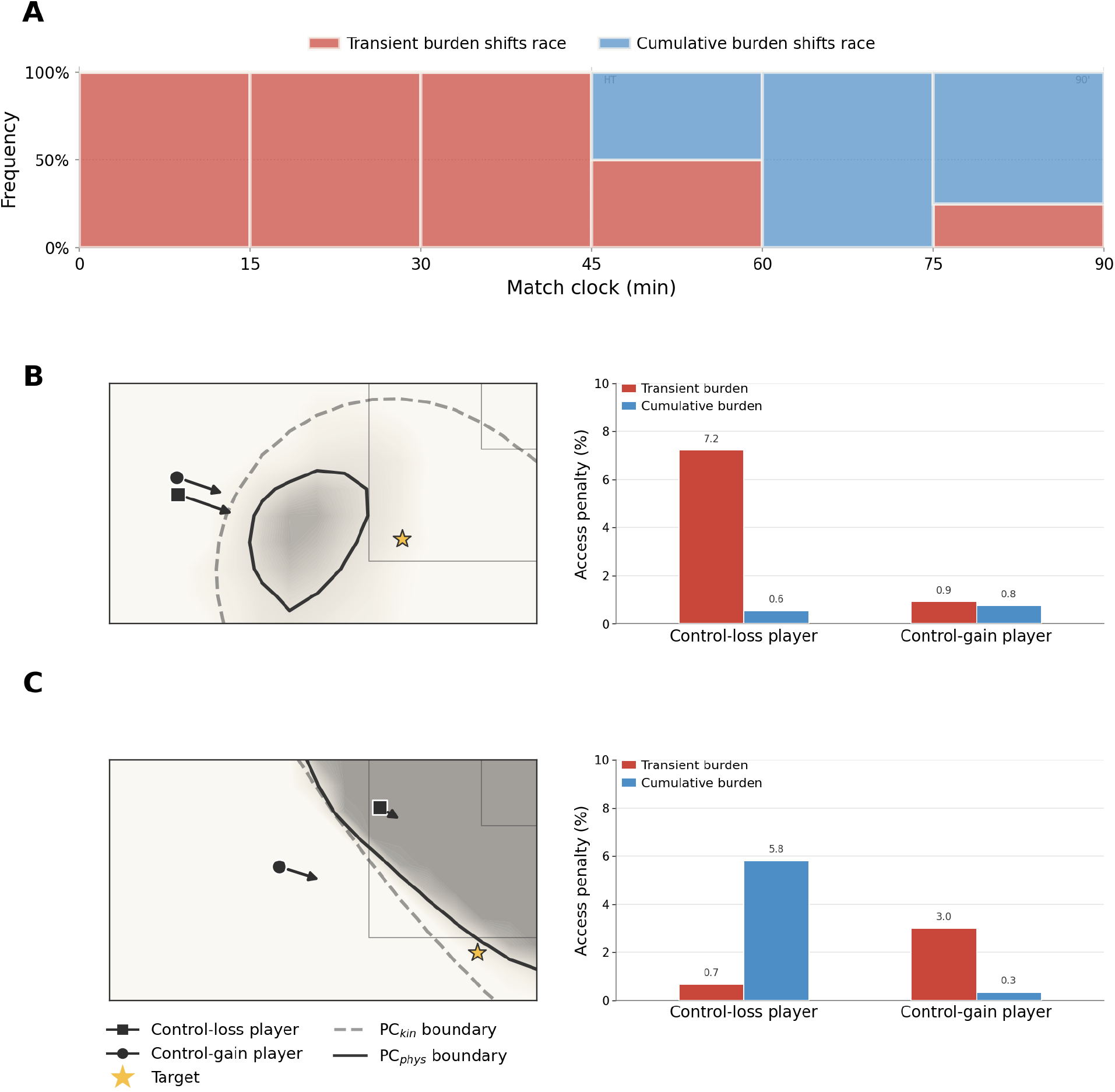
Head-to-head race application. **A:** Tournament-wide composition of physiology-driven race shifts by match clock (15-minute bins). Red = transient-dominant (*Z*); blue = cumulative-dominant (*G*). Transient-driven shifts are more frequent in the first half; cumulative-driven shifts account for the majority from approximately the 60th minute onward. **B:** Transient-dominant exemplar (Argentina vs France, WC Final, 11:40). Varane (France centre-back) is the control-loss player, carrying a 7.2% transient access penalty against 0.6% cumulative as Alvarez (Argentina centre-forward) attacks the target space; the Phys-PC boundary (solid) contracts relative to the kinematic boundary (dashed). **C:** Cumulative-dominant exemplar (France vs Denmark, 88:12). The access penalty is predominantly cumulative (5.8%, *G* channel) with minimal transient burden.

Figure 6B shows a transient-dominant case (Argentina vs France, WC Final, 11:40): Varane, a France centre-back, carries a 7.2% transient access penalty against only 0.6% cumulative, while Alvarez, an Argentina centre-forward, is physiologically unencumbered in his approach. The Phys-PC model identifies Varane as the more constrained contestant in this race. What followed illustrates a broader pattern: a second French defender intervened to physically narrow Alvarez’s approach angle, deploying a collective tactical adjustment that a reachability surface cannot capture. The exemplar is therefore informative not because the outcome confirmed the model, but because it shows how a transient physiological gap can create a structural vulnerability that the team must address through means other than the primary contestant’s individual recovery.

Figure 6C shows a cumulative-dominant case (France vs Denmark, 88:12): Norgaard, a late substitute with near-zero *G* accumulation, contests the ball against Upamecano, a starter carrying 5.8% cumulative capacity loss. The kinematic model assigns the access advantage to Upamecano, while physTTA reverses this assessment based on the G differential. Norgaard won the subsequent contest; this outcome is consistent with the reserve differential that Phys-PC identified, though the model does not claim to predict individual contest results from physiological state alone.

### 6.2 Transition and counterattack vulnerability

#### 6.2.1 Motivation

Ball-winning transitions out of possession-dominant phases test both physiological channels simultaneously, but in different players. During sustained attacking possession the effort distribution is not uniform: wingers and attacking midfielders performing explosive runs and final-third 1v1s accumulate substantial transient burden, especially compared to central defenders cycling the ball at lower intensity in stable positions further back. When possession is lost, both groups must recover, but their physiological states at that moment reflect what they were doing in the seconds and minutes before, not just how long they have been on the pitch. A reachability model that assumes constant capacity across all players will fail to capture this asymmetry between team lines.

#### 6.2.2 Event extraction

Transition candidates are possession changes in open play where, at the turnover frame, the ball is at least 5 m inside the new attacking team’s own half and both outfield team centres of mass are also in that half. The candidate must develop into a meaningful counterattack: the ball advances at least 20 m in the attacking direction, crosses the halfway line, and the new team maintains continuous possession for at least 5 seconds without a stoppage or further possession change. Set-piece restarts are excluded.

#### 6.2.3 Findings

At the transition release frame, Phys-PC computes the full-pitch TTA surface for all outfield players. The difference between the Phys-PC and kinematic pitch-control surfaces for the (now defending) team identifies regions where physiological state reduces model-assessed coverage below the kinematic baseline.

The role-by-zone heatmap (Figure 7A) reveals a pattern that maps directly onto the expected activity profile of each position during sustained possession. Wingers and attacking midfielders operating in the final third are predominantly transient-dominant at the transition moment: these players are most engaged in explosive 1v1 efforts, forward runs, and high pressing, all of which accumulate transient burden rapidly. Central defenders, by contrast, are characteristically cumulative-dominant as they hold the defensive shape and provide stable recycling outlets during possession. Midfielders occupy an intermediate zone, reflecting their dual role as connectors between phases. The role-depth structure in Figure 7A is therefore not incidental; it is a direct readout of which players accumulated which type of burden by doing their job in the possession phase immediately before the transition.

**Figure 7.**
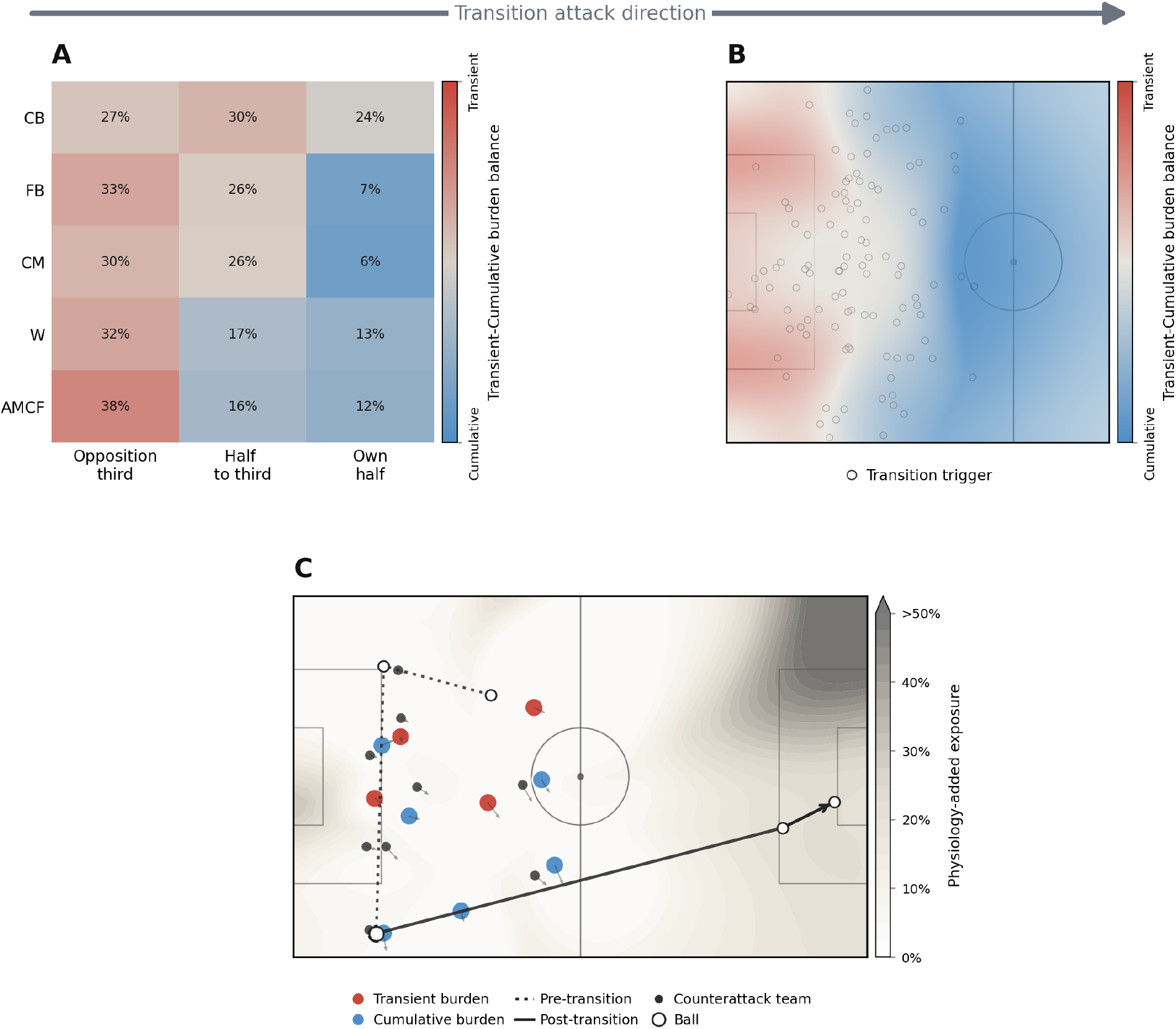
Transition and counterattack vulnerability application. **A:** Role-by-zone burden-balance heatmap across all tournament transitions at the possession switch. Red = transient-dominant (*Z*); blue = cumulative-dominant (*G*). Attacking midfielders and forwards pressing in the opposition third are predominantly transient-dominant; fullbacks and central defenders in their own half are more cumulative-dominant. **B:** Spatial aggregate of the transient-cumulative burden balance at all transition trigger locations (circles). Transient dominance concentrates near the pressing contest; cumulative dominance characterises deeper recovery positions. **C:** Exemplar transition (France vs Denmark, 90:11). All players are shown; Denmark players, now defending after the turnover, are coloured by dominant burden channel, while France counterattacking players are shown as small dark dots. The ball trajectory (black line) runs from own half to the final third. The grey Phys-PC exposure surface shows where the defending team’s model-assessed coverage falls below the kinematic baseline.

The spatial aggregate (Figure 7B) translates this into pitch geography. The half-spaces in the final third where wingers and attacking midfielders are most active during possession are characterised by high transient burden at the transition moment. The deeper band corresponding to the defensive midfield and central defenders shows greater cumulative burden dominance. These are not equivalent constraints for recovery runs. Transient-burdened players in the final third face the longest recovery distances with the highest capacity penalty, creating structural rifts between the pressing line and the defensive block. The players who expended energy creating openings are the same players who must now bridge the gap to the central defenders, and they are the most physiologically constrained to do so. Kinematic pitch control, which sees none of this, assigns each player the same capacity regardless of their pre-transition role.

The exemplar (Figure 7C; France vs Denmark, 90:11) shows this in action where Denmark lost the ball to France, who immediately transitioned into a counterattack. Players involved in the pre-transition buildup for Denmark (i.e., those near the ball when possession was lost) appear predominantly red (transient-burdened), reflecting the load from their involvement in the attacking sequence. Players further from the ball at the moment of the switch are predominantly blue (cumulative-burdened), reflecting accumulated match load. The Phys-PC exposure surface concentrates the largest coverage reductions precisely in the counterattack corridor where the transient-burdened players would need to recover. Kinematic pitch control underestimates vulnerability in these transitions overall, but most acutely for the players whose transient state renders their recovery capacity most constrained at that moment.

A concrete illustration of why this matters: at the transition frame, Denmark had four players near the halfway line against two French attackers advancing on the counterattack, indicating a numerical defensive advantage that a kinematic model reads as ample coverage. Two of those four Danish players, however, carried high transient burden from their involvement in the preceding attacking sequence, as reflected in their dominant state (red: transient burden). The effective recovery contest was therefore closer to a 2v2 than the raw headcount suggests, thus illustrating a vulnerability that kinematic pitch control, treating all four players as equivalently capable, does not reveal.

### 6.3 Player and role profiling by valuable-space retention

#### 6.3.1 Motivation

The previous applications focus on event-level reachability. A complementary use case is profiling: identifying how players and role groups maintain access to valuable territory across match load. The question is not simply which player reaches the most space, but whether that access is carried through repeated high-intensity actions or through sustained positioning and accumulated involvement. This reframes Phys-PC from a single-event adjustment into a diagnostic of how valuable-space access is physiologically supported.

#### 6.3.2 Metric construction

For each player, we compute actor-isolated kinematic and physiological pitch-control surfaces and integrate them over a value-and-density map that upweights goal-facing, ahead-of-ball, defensively contested territory. The horizontal axis in Figure 8 is the kinematic contested space access 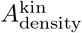 (m^2^ per 90 minutes): how much contested, goal-relevant territory a player can reach from their current position and movement state, before any physiological penalty is applied. Higher values indicate larger actor-isolated access to contested space, not necessarily better performance.

**Figure 8.**
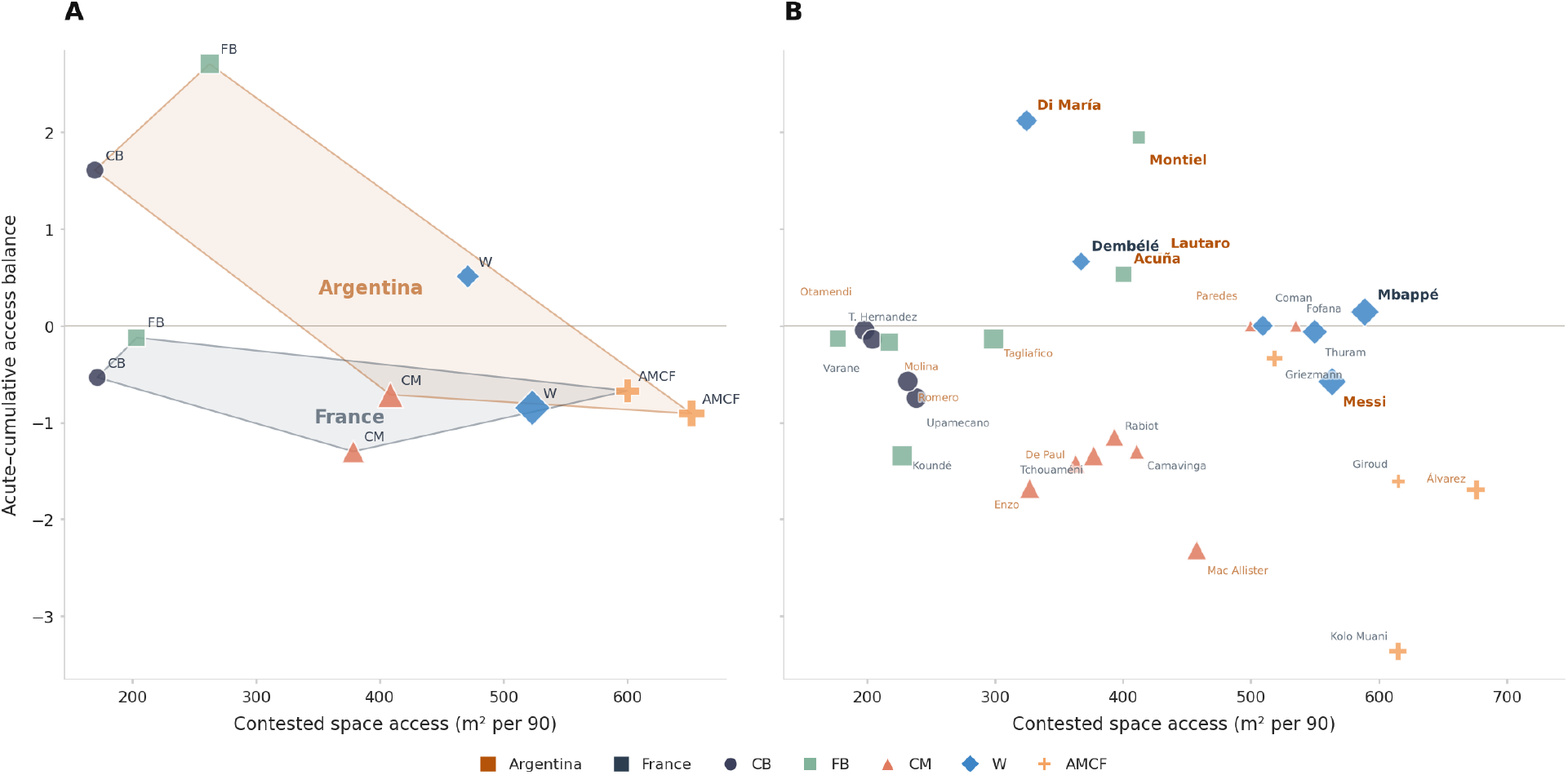
Player-profiling application. **A:** Tournament-wide France and Argentina team-role aggregates, ranked against other teams’ same-role aggregates. The x-axis is contested space access (m^2^ per 90): the kinematic actor-isolated pitch-control integral over a value-and-density surface. The y-axis is the acute–cumulative access balance, computed as the log ratio of role-conditional percentile ranks of acute and cumulative fatigue cost. Positive values indicate acute burst-recovery dominance; negative values indicate cumulative-load dominance. Shaded hulls enclose each team’s role points; bubble size scales role share of team contested-space volume; colours and markers denote positional role. **B:** Player-level profiles from the France–Argentina final, restricted to players with at least 15 minutes on pitch. The y-axis is ranked against the France+Argentina tournament player-match pool within the same role. Player labels are coloured by team. Bubble size scales minutes played.

The vertical axis captures the *mode* of physiological erosion. Total fatigue cost is 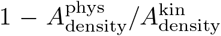; this is decomposed into a cumulative component (the shortfall with only the cumulative *G* channel) and an acute residual. We then plot the log ratio of the role-conditional percentile rank of acute cost to cumulative cost, so that positive values indicate access whose physiological cost is dominated by recent repeat-action demands (sprints, overlaps, pressing), while negative values indicate access shaped more by sustained role occupation and accumulated match involvement. Role conditioning ensures that a winger is compared against other wingers, not against centre-backs. Because these profiles are computed from actor-isolated surfaces, they describe the physiological mode through which each player retains their own valuable-space access, not whether teammates compensate for any reduction.

#### 6.3.3 Findings

The team-role profiles first show a clear positional stratification along the horizontal axis (Figure 8A). Centre-backs and fullbacks occupy lower contested-access values, central midfielders sit in the middle of the range, and winger/attacking-midfield-forward roles carry the largest actor-isolated access to contested territory. This ordering is not a player-quality ranking. It reflects where the value-and-density surface assigns opportunity: advanced and connected roles are more often positioned to threaten goal-facing, ahead-of-ball, contested space, while deeper defensive roles access that territory less frequently in actor-isolated terms.

The vertical axis adds a different layer. For both teams, central midfield and attacking roles lie on the sustained-involvement side of the profile space. This is consistent with positional access: these roles create value not only through isolated bursts, but by repeatedly being available between lines, around the ball, or in forward receiving positions. A central midfielder can retain access to valuable passing and support spaces by staying connected to play; an attacking midfielder or forward can retain access to advanced territory by holding positions that keep short paths into dangerous areas. Their physiological erosion is therefore more closely associated with accumulated involvement than with a recent burst-recovery episode. Put differently, the model separates a role’s territorial volume from the way that volume is physiologically maintained.

The team contrast is then more informative than a simple above/below-zero reading. France lies below the role-balanced line across all five role groups. This should not be read as an absence of pace. France could still threaten space explosively, but their role-level access to valuable territory was shaped more by sustained positioning and accumulated match involvement than by repeated recent burst demands. In practical terms, a French forward or winger could threaten depth because their starting position already preserved access to dangerous space, rather than because the role repeatedly had to leave that space and sprint back into it. The profile therefore suggests a relatively homogeneous positional-access signature: across roles, France’s valuable-space access was not primarily characterised by acute repeat-action debt, but by players repeatedly occupying positions from which valuable space was already within reach.

Argentina shows a more polarised role structure. Its central and attacking-platform roles share the sustained-involvement profile, but centre-backs, fullbacks, and wingers shift to the repeat-action side. The fullback group is the strongest acute-dominant team-role point among the finalists, suggesting that valuable flank access was more directly tied to overlapping, recovering, and re-accelerating actions. In this profile, access to valuable wide territory is not simply held by starting position; it is more often rebuilt through recent movement demands. The centre-back signal should be interpreted similarly cautiously: not as high territorial volume, but as evidence that the valuable access they did provide was more coupled to recent stepping, covering, or recovery demands. The contrast with France is therefore tactical: France’s profile is more consistent with positionally held access, stable spacing, or more selective timing of explosive actions, whereas Argentina’s profile suggests that wide and defensive-wide access was more action-generated and carried more of the repeat-action burden.

At the player level (Figure 8B), individual profiles redistribute the team-role signatures in revealing ways. Di Maria, Dembele, and Lautaro Martinez occupy a lower-volume, acute-skewed region, consistent with episodic, action-generated access: players entering valuable spaces through timed bursts rather than continuously holding a broad platform. Mbappe represents a different departure from the French winger norm: his acute shift does not coincide with reduced contested-access volume. He retains high-access status, but his final-match profile moves above France’s sustained-access winger signature, combining sustained high-value territorial presence with acute burst-mediated entry into that territory.

Argentina’s final profile separates the wide channel into three physiological modes. Di Maria supplies the selective acute attacking spike; Messi, though labelled as a winger, sits below Argentina’s acute wide-role tendency, closer to a sustained-access playmaker profile in which valuable territory is preserved through positioning and connection to possession; and the fullbacks (Montiel and Acuna) carry the larger wide platform at above-norm contested-access volume, with Montiel also strongly acute-skewed. Argentina’s wide access in the final was therefore not simply winger-driven but distributed across three complementary physiological signatures.

## 7 Discussion

The preceding sections establish that physiology-aware pitch control produces state dynamics consistent with exercise-physiology benchmarks, associates with match outcomes at three tactical scales, and reveals spatial structure that kinematic models cannot detect. We now consider the magnitude of these effects, the methodological boundaries of the approach, and how Phys-PC relates to the broader pitch-control literature.

Across the three outcome families, the consistent signal is not that high reserve alone guarantees success, but that *relative physiological asymmetry* helps explain which side can convert space into an observed football outcome. The validation odds ratios (1.20-1.45) are modest when considered as standalone predictors, but this framing understates their role. Physiological reserve is one input to a contest that is simultaneously shaped by technique, positioning, anticipation, and tactical instruction. A 4-8 percentage-point adjusted difference in success rate is small relative to the tactical and technical variability present in any single action. However, effects of this magnitude may accumulate across repeated contests over the course of a match, making physiological asymmetries relevant even when no individual event is strongly determined by physiological state alone. The effects are directionally aligned across individual contests, race-to-space passing, and team-level territorial progression, and they survive event-geometry, match-context, and random-intercept controls. Phys-PC is therefore not primarily a standalone pass-completion or duel-outcome predictor; it is a relative advantage metric that quantifies where physiological asymmetries exist between opponents and which tactical actions appear to benefit from, or be harmed by, those asymmetries.

A methodological caveat follows directly from this framing. Physiological state is inferred from the same spatiotemporal tracking data used to compute kinematic TTA, and the validation outcomes (duel wins, pass completions, final-third entries) are themselves determined by many non-physiological factors. The associations reported here therefore demonstrate consistency between inferred physiological asymmetry and observed outcomes, not causal confirmation. The model identifies when a relative physiological gap exists between opponents before an action; it does not establish that the gap caused the outcome. This is a deliberate interpretive boundary. External validation against directly measured internal-load indices (heart rate, blood lactate, neuromuscular markers) or substitution-timing data would be needed to strengthen causal claims about the inferred penalties.

A related limitation is the model’s inability to distinguish physiological constraint from deliberate pacing. A player who could sprint but chooses not to, whether for energy conservation or positional discipline, is indistinguishable in the current framework from a player whose transient or cumulative physiological state genuinely limits the effort. This conflation is inherent to any tracking-only fatigue inference and means that Phys-PC estimates physiological affordability rather than behavioural intent Noakes [2000]. The practical consequence is that some fraction of the access reduction attributed to physiological burden may instead reflect tactical choice, particularly in roles where pacing is strategically managed (e.g., centre-forwards conserving effort between pressing triggers).

Phys-PC is situated within a growing line of work that extends pitch-control models beyond uniform kinematic assumptions. Da Silva et al. da Silva et al. [2025] demonstrated that replacing a uniform top speed with player-specific *v*_95_ values produces a measurable shift in team-level pitch control, with the magnitude depending on positional role and match phase. Their stamina factor *ξ* modulates *v*_max_ by a scalar multiplier, revealing a logarithmic relationship between speed scaling and aggregate pitch control. Phys-PC shares the premise that kinematic homogeneity is insufficient, but decomposes the scalar adjustment into two mechanistically grounded channels with distinct temporal dynamics: a transient channel that fluctuates on the timescale of sprint-recovery cycles and a cumulative channel that evolves monotonically across the match. This decomposition enables action-level inference (which channel dominates a specific race or transition) rather than match-aggregate description. The player-profiling application extends this perspective by characterising not only how much valuable territory a player can reach, but also the physiological mode through which that access is sustained. This distinction separates access outcomes from the mechanisms that produce them, analogous to prior work that distinguished between occupying space and generating it through off-ball movement Fernandez and Bornn [2018].

### 7.1 Limitations and future work

Several simplifying assumptions bound the interpretation of these results. The movement model computes physiological demand along the direct ray from a player to each target location. This preserves compatibility with standard pitch-control formulations, but it does not explicitly model curved tactical paths, evasive routes, contact-mediated movement costs, or the full mechanical demands of change-of-direction actions. The alignment penalty partially captures directional cost, but remains a surrogate for the deceleration, re-acceleration, and eccentric loading associated with high-COD movements Yamashita et al. [2025], Proske and Morgan [2001]. Similarly, the metabolic-power proxy should be interpreted as an estimate of locomotor demand during accelerated running and is less reliable for shuffling, curvilinear motion, braking-dominant actions, and contact situations Hader et al. [2016].

The physiological state variables are also simplified representations rather than directly measured quantities. *Z* summarizes rapidly reversible burden, *G* represents cumulative match-scale drain, and the access scale *ρ* estimates reduced physiological availability rather than a specific internal-load metric. In the present implementation, calibration is performed at the position-role level and relies on literature-informed parameterization, limiting the ability to capture player-specific differences in sprint capacity, recovery kinetics, and pacing behavior. The related limitation that the model cannot distinguish physiological constraint from deliberate pacing is discussed above.

Future work should improve each layer of the model while preserving its modular structure. Route-planning or Eikonal formulations could replace the direct-ray assumption and incorporate curved paths, opponent-shaped corridors, and explicit change-of-direction costs. Player-specific recovery and power-duration parameters could be estimated using hierarchical models calibrated against longitudinal tracking, heart rate, RPE, or other internal-load measures. Probabilistic extensions could propagate uncertainty in physiological access through the pitch-control surface, while external validation against substitution timing, sprint decrement, neuromuscular markers, or club internal-load indices would strengthen confidence in the magnitude and practical interpretation of the inferred penalties.

## 8 Conclusion

Pitch control models have treated spatial reachability as a function of kinematics alone. Phys-PC extends this to ask a prior question: given what this player has done over the course of the match and in the moments just before this action, how much of their kinematic capacity remains available? The answer depends on both the transient recoverable burden from recent high-intensity efforts, and the cumulative non-recoverable drain from match-scale expenditure above the aerobic threshold. Practical match examples show that these channels leave distinct signatures in individual races, team-level spatial vulnerability at transition moments, and player-role profiles of valuable-territory retention. By operating as a TTA pre-processor calibrated entirely from broadcast tracking data, Phys-PC integrates into any existing pitch-control framework without modifying the downstream spatial model and without requiring laboratory measurements or club-collected internal load data.

## Code availability

The code generated during this study is publicly available on GitHub at https://github.com/abdulzaf/phys-pc. A version-tagged, citable snapshot corresponding to this manuscript is archived on Zenodo at https://doi.org/10.5281/zenodo.21197841.

## Acknowledgements

The authors have no acknowledgements to declare.

## A Supplementary robustness analysis

To assess whether the outcome-facing conclusions depend on a narrow parameter setting, we performed a one-at-a-time sensitivity analysis around the frozen manuscript parameters entering the reserve-access scale. The following quantities were perturbed by ±10% and ±20%: cumulative burden weight, transient burden weight, transient activation threshold, transient recovery time constant, *G*-drain magnitude, and minimum capacity/access floor. The access weights, activation threshold, and floor were recomputed directly in the access-scale equation. Because the exported outcome samples contain post-cache physiological states rather than the full pre-event power history, the transient recovery and *G*-drain checks were implemented as transparent state-level proxies: transient recovery was represented by scaling the exported *Z* state, and *G*-drain magnitude by scaling capacity loss 1 − *φ*(*G*). The movement-alignment penalty was not included in this outcome-facing check because the Section 5 outcome models use *ρ* gaps rather than directional attenuation.

For each perturbation, the outcome analyses reused the frozen event samples and recomputed the relevant relative reserve gap before assigning quartiles. We then calculated the raw Q4-versus-Q1 odds ratio and a univariate quartile trend odds ratio. These robustness endpoints are intentionally simpler than the main GLMMs: they test whether the direction of the physiological-asymmetry signal is stable under modest parameter changes, without re-tuning the model or introducing mixed-model convergence as an additional source of variation.

The outcome-facing results remain directionally positive across all access-scale perturbations: the highest relative-reserve quartile retains higher raw odds than the lowest quartile for ground challenges, over-the-top passes, and middle-third possession sequences. These checks support the interpretation that the Section 5 conclusions are not artifacts of a single brittle parameter choice, while leaving full individual-parameter uncertainty estimation to future work with richer internal-load data.

**Table 5:** Sensitivity summary for one-at-a-time parameter perturbations. Ranges are the minimum and maximum values across the baseline and all access-scale perturbations for the relevant endpoint. Rows report raw Q4-versus-Q1 odds ratios on the frozen Section 5 event samples.

| Endpoint | Baseline | Range | Stable variants |
| --- | --- | --- | --- |
| Ground challenges: Q4 vs Q1 OR | 1.14 | 1.06-1.21 | 25/25 |
| Over-the-top passes: Q4 vs Q1 OR | 1.35 | 1.03-1.36 | 25/25 |
| Possession sequences: Q4 vs Q1 OR | 1.26 | 1.19-1.34 | 25/25 |

